# Powering single-cell analyses in the browser with WebAssembly

**DOI:** 10.1101/2022.03.02.482701

**Authors:** Aaron Lun, Jayaram Kancherla

**Author notes:** Authors contributed equally to this work.

## Abstract

We present kana, a web application for interactive single-cell ‘omics data analysis in the browser. Like, literally, in the browser: kana leverages web technologies such as WebAssembly to efficiently perform the relevant computations on the user’s machine, avoiding the need to provision and maintain a backend service. The application provides a streamlined one-click workflow for the main steps in a typical single-cell analysis, starting from a count matrix and finishing with marker detection. Results are presented in an intuitive web interface for further exploration and iterative analysis. Testing on public datasets shows that kana can analyze over 100,000 cells within 5 minutes on a typical laptop.

## 1 Background

Modern web browsers are sophisticated pieces of software responsible for rendering, scripting, networking, data storage and more. New web technologies such as WebAssembly [23] have greatly enhanced browsers’ capabilities for intensive computation, providing new opportunities to repurpose the browser as an interactive data analysis tool. The idea is to use the browser itself to perform statistical and computational analyses directly on the user’s machine, i.e., “client-side compute”. Those analysis results can then be rendered on a web page for convenient visualization and exploration within a familiar interface.

This paradigm of client-side compute in the browser has several advantages over the traditional model of server-side data analysis. Most obviously, we do not need a scalable backend server to perform any of the calculations, simplifying deployment and reducing costs. As the entire analysis is performed on the client machine, we avoid any data transfer - this reduces latency in the user interface and circumvents concerns over data ownership and privacy. Finally, we do not require installation of any data analysis environments like R or Python, ensuring that the analyses are accessible to audiences of varying computational skill.

Some existing web applications have successfully realized the benefits of client-side compute for bioinformatics data analysis. The BioJS registry hosts JavaScript components for handling biological data [19], many of which focus on DNA and protein sequence analysis. The BrowserGenome.org application performs read alignment and transcript quantification for RNA sequencing data [60], while the ubit2 application analyzes single-cell quantitative polymerase chain reaction datasets [18]; both use JavaScript implementations of the relevant algorithms to run in the browser. However, these are exceptions to the rule, and most bioinformatics web applications only use the browser to visualize results that were computed elsewhere, e.g., cellxgene [54], Epiviz [13], Cirrocumulus [20], Vitessce [27], HiGlass [28] and every R/Shiny application ever [12].

The client-side paradigm holds great potential for single-cell genomics due to the exploratory nature of its data analysis. Most use cases for single-cell sequencing involve identifying new cell subpopulations or states from heterogeneous biological samples [62], which requires several iterations of data analysis, visualization and interpretation. If the compute could be performed in the browser, the results could be immediately rendered on a web page for a seamless transition between analysis and interactive exploration. Of course, this is complicated by the size of the datasets, each of which involves tens of thousands of genes and up to millions of cells; and the lack of browser-compatible implementations of various algorithms, most of which are written in R or Python.

We present kana, a web application for analyzing single-cell ‘omics data inside the browser. kana provides a streamlined one-click workflow for the main steps in a typical single-cell analysis [2], starting from a count matrix (or multiple such matrices, for datasets with multiple modalities and/or samples) and finishing with marker detection. Users can interactively explore the low-dimensional embeddings, clusterings and marker genes in an intuitive graphical interface that encourages iterative re-analysis. Once finished, users can save their analysis and results for later examination or sharing with collaborators. By using technologies like WebAssembly and web workers, we achieve high-performance compute for datasets containing hundreds of thousands of cells.

The kana application is available at https://www.kanaverse.org/kana. Developers can set up their own deployments by following the instructions at https://github.com/kanaverse/kana.

### 2 Results

### 2.1 Application overview

Given a single-cell dataset - typically for gene expression, but possibly with protein and/or CRISPR counts - kana implements a routine analysis with the steps listed below. We will not discuss the statistical and scientific rationale behind each step in much detail as this has been covered elsewhere [44].

1. We import a feature-by-cell count matrix from the user’s machine. This can take the form of Matrix Market files such as those produced by the Cellranger pipeline; HDF5 files, using either the 10X HDF5 feature barcode matrix format or as H5AD files; SummarizedExperiment or SingleCellExperiment objects [2] saved to RDS files; or datasets from public repositories like Bioconductor’s ExperimentHub [56].
2. We compute common quality control (QC) metrics such as the total count, number of detected genes and proportion of mitochondrial counts. Low-quality cells are defined as those cells with outlier values for any of these metrics; these are filtered out from subsequent steps.
3. We perform scaling normalization based on the library size to remove cell-specific biases. This is followed by a log-transformation to obtain a matrix of log-normalized expression values. For ADT counts, this uses a grouping approach to reduce the impact of composition biases [32].
4. We fit a LOWESS trend [14] to the per-gene variances with respect to the means, computed from the log-expression values. We sort on the residuals to define a subset of highly variable genes (HVGs).
5. We perform principal components analysis (PCA) on the log-expression matrix with the subset of HVGs. This yields a few top PCs that capture the heterogeneity of the data in a compressed and denoised form.
6. For datasets containing multiple batches, we remove effects between cells from different samples using mutual nearest neighbors (MNN) correction [24]. This can also be used to integrate multiple datasets based on their set of shared features.
7. We apply clustering techniques on the top PCs to generate discrete sub-populations. By default, this uses various flavors of community detection on a shared nearest neighbor (SNN) graph where each cell is a node and edges connect neighboring cells. We also support k-means clustering with some user-specified choice of the number of clusters.
8. Each cluster is characterized through differential expression analyses to detect its marker genes. Specifically, we perform a series of pairwise comparisons between clusters and summarize the effect sizes into a ranking of potential marker genes for each cluster. Users can also directly compare two clusters of interest to characterize differences between two closely related groups of cells.
9. We perform gene set enrichment analyses on the top markers for each cluster with publicly available collections like the Gene Ontology [4]. We can also compute and visualize per-cell scores for individual gene sets using a PCA-based method [11].
10. We perform further dimensionality reduction on the top PCs to obtain two-dimensional embeddings for visualization. This includes the usual t-distributed stochastic neighbor embedding (t-SNE) and uniform manifold approximation and projection (UMAP) [65, 53].
11. We can optionally annotate cell types for each computed cluster using the SingleR algorithm [3]. This is based on labelled reference datasets involving a variety of human and mouse cell types.
12. For CITE-seq [63] and Perturb-seq datasets [16], we can combine information from multiple modalities during clustering and visualization. This uses a nearest-neighbor approach to equalize the baseline variance of each modality, though user-specified weights are also supported. Markers are also computed separately for each modality.

At each step, users can easily customize key parameters (Figure 1). For example, we can adjust the QC thresholds, the number of HVGs and top PCs, the granularity of the clustering, and more. Iterative refinements to the parameters are encouraged in kana, as the application tracks dependencies between steps to enable efficient re-analysis. Specifically, when parameters are modified for any step, all subsequent steps are automatically re-executed to propagate the change to downstream results. Conversely, kana does not rerun any steps upstream of the change to avoid unnecessary recomputation and reduce latency.

**Figure 1:**
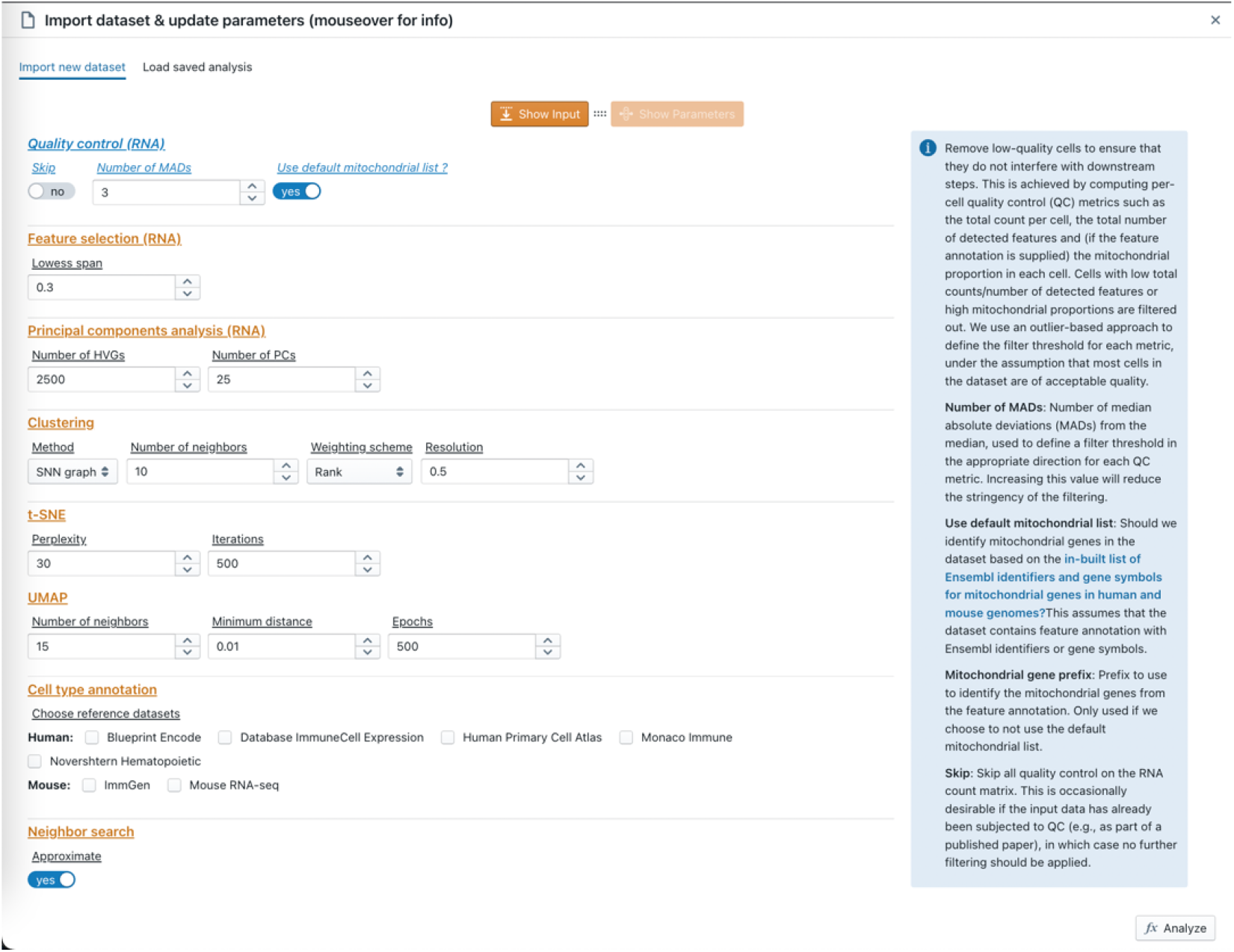
Screenshot showing the analysis configuration panel in the kana application. Clicking “Analyze” will perform the entire analysis.

Once each step of the analysis is complete, kana visualizes its results in a multi-panel layout (Figure 2). One panel contains a scatter plot for the low-dimensional embeddings, where each cell is a point that is colored by cluster identity or gene expression. Another panel contains a table of marker statistics for a selected cluster, where potential marker genes are ranked and filtered according to the magnitude of upregulation over other clusters. We also provide a gallery to visualize miscellaneous details such as the distribution of QC metrics. Finally, users can export the analysis configuration and results for later inspection. The exported results can be quickly reloaded in a new browser session, allowing users (or their collaborators) to explore those results without repeating the computation. Similarly, the exported configuration can be used to restore the analysis session for further iteration.

**Figure 2:**
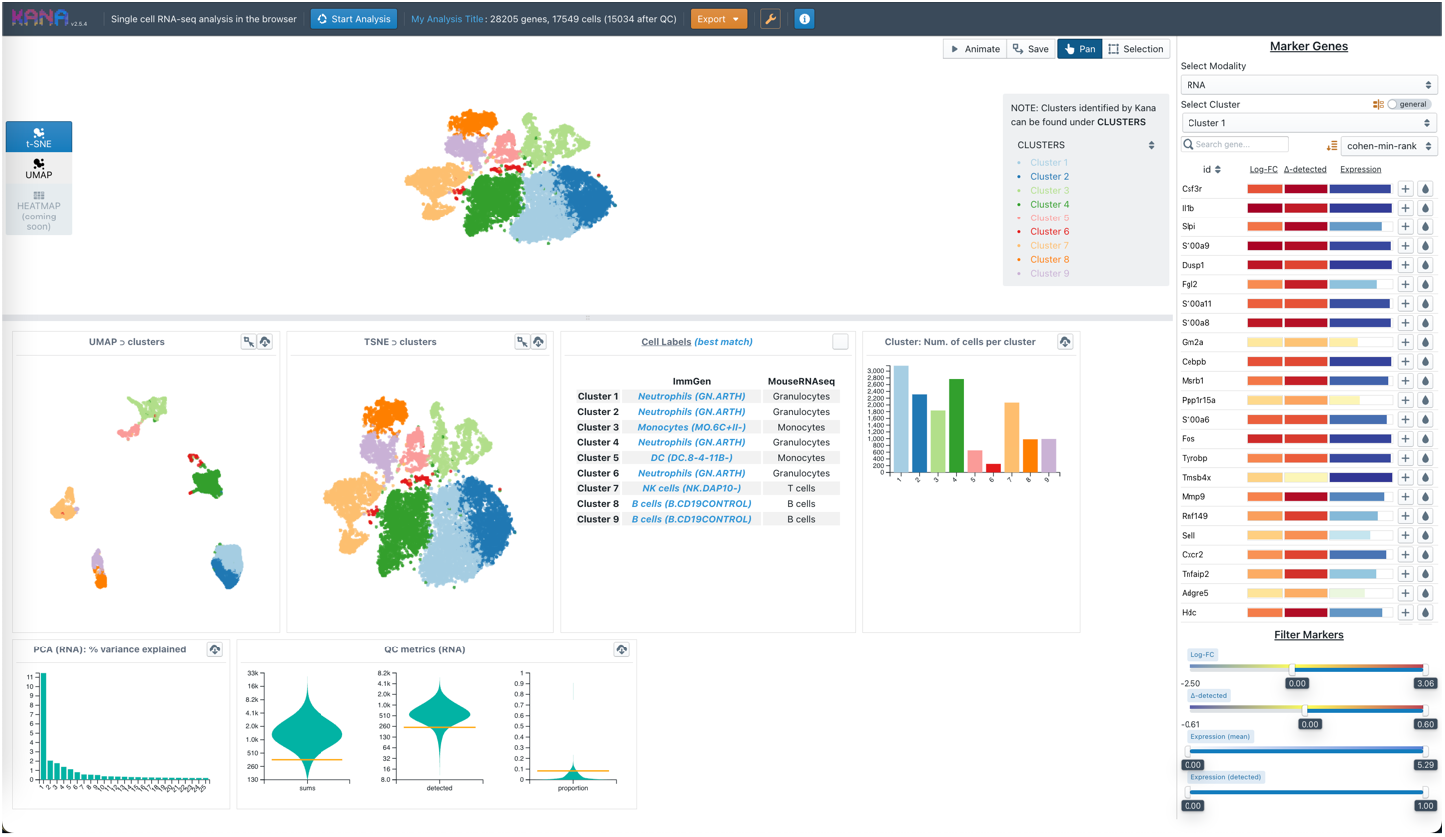
Screenshot showing the multi-panel layout for results in the kana application. The top-left panel is used for the low-dimensional embeddings, the right panel contains the marker table for a selected cluster, and the bottom-left panel contains a gallery of miscellaneous plots.

### 2.2 Client-side computation

#### 2.2.1 Efficient compute with WebAssembly

WebAssembly (Wasm) [23] is a binary instruction format that provides a web-executable compilation target for languages like C/C++, Go and Rust. It aims to provide near-native performance for computationally intensive tasks, running alongside and complementing JavaScript in web applications. The use of Wasm allows us to convert the browser into a compute engine by integrating existing scientific libraries for bioinformatics data analysis - see the biowasm project [1] for previous efforts in this direction. For kana, we collected or created C++ implementations of the algorithms required for each analysis step:

- The tatami library [46] provides an abstract interface to different matrix classes, based on similar ideas in the beachmat package [45]. In addition to the usual dense and sparse representations, tatami also supports the delayed operations implemented in the DelayedArray package [22]. This allows the creation of QC-filtered and log-normalized matrices without any duplication of expression data.
- The CppWeightedLowess library [40] contains a C++ port of the weightedLowess function from the limma package [59]. This function is, in turn, based on the LOWESS algorithm [14] implemented in R’s lowess function, modified with the ability to consider weights in the span calculations.
- The CppIrlba library [34] library contains a C++ port of the IRLBA algorithm [8] to efficiently obtain the top PCs from an input matrix. This is based on the C code in the irlba R package [9] with some refactoring to eliminate R-specific dependencies. In particular, we now rely on the Eigen library [21] for matrix operations.
- The CppKmeans library [35] implements the Hartigan-Wong [25] and Lloyd algorithms [31] for k-means clustering. The Hartigan-Wong implementation was translated from the Fortran code used by R’s kmeans function. We also provide several initialization methods including kmeans++ [66] and PCA partitioning [64].
- The knncolle library [39] wraps several nearest neighbor detection algorithms in a consistent interface. This includes exact methods like vantage point tree search [67] as well as approximate methods like Annoy [10]. Its design is based on the BiocNeighbors package from Bioconductor [33].
- The libscran library [36] implements high-level methods for single-cell RNA sequencing (scRNA-seq) data analysis, ranging from quality control to clustering. The code here originates from the scran, scuttle and scater packages [51, 52] bundled together into a single C++ library for convenience. This library relies on the igraph C library [15] for community detection from the SNN graph.
- The qdtsne library [37] contains a C++ implementation of the Barnes-Hut t-SNE algorithm [65]. This is mostly a refactored version of the code in the Rtsne package [29]. Some additional optimizations have been applied to improve scalability.
- The umappp library [38] contains a C++ implementation of the UMAP algorithm [53]. This is derived from code in the uwot R package [55].
- The SinglePP library [43] contains a C++ implementation of the SingleR algorithm [3]. This is derived from code in the SingleR Bioconductor package.
- The CppMnnCorrect library [41] implements the MNN method fo batch correction [24]. It is loosely based on the fastMNN method from the batchelor Bioconductor package [30].
- The rds2cpp library [42] implements an RDS file parser/writer as a standalone C++ library. This allows kana to read datasets from RDS files without linking to R itself.

We created the scran.js library [50] to provide JavaScript bindings to these C++ libraries. Specifically, we compiled our C++ code to Wasm using the Emscripten toolchain [68], allowing applications to perform single-cell-related calculations from JavaScript by calling scran.js functions. The same libraries can also be used in other web applications via a standard NPM installation, or outside the browser if a suitable Wasm runtime environment is available. For example, the bakana library wraps scran.js functions into an analysis pipeline for “one-click” execution in both the browser and Node.js environments.

To evaluate the efficiency of our Wasm strategy, we compared a kana analysis in the browser to that of a native executable compiled from the same C++ libraries [47]. We analyzed several public scRNA-seq datasets (Table 1) using the default kana parameters for both approaches, i.e., QC filtering to 3 MADs from the median; PCA on the top 2500 HVGs to obtain the top 25 PCs; SNN graph construction with 10 neighbors and multi-level community detection at a resolution of 0.5; t-SNE with a perplexity of 30; UMAP with 15 neighbors and a minimum distance of 0.01; and 8 threads for all parallel sections (i.e., web workers for kana, see below). We collected timings on an Intel Core i7-8850H CPU (2.60GHz, 6 cores, 32 GB memory) running Manjaro Linux. For convenience, we ran the kana timings in batch using Puppeteer [5] to control a headless Chrome browser (HeadlessChrome/98.0.4758.0).

**Table 1:**
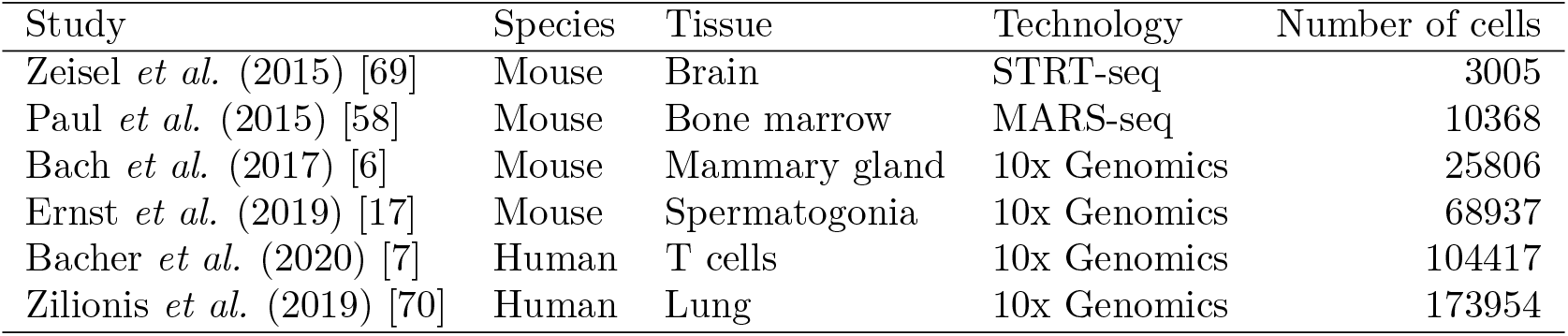
Collection of scRNA-seq datasets used for testing.

Our results indicate that kana analyses took approximately 25-50% longer to run compared to the native executable (Table 2). This is consistent with other benchmarking results [26] where the performance gap is attributed to Wasm’s design constraints and the overhead of the browser’s Wasm runtime environment. Our native executable was also created with a different compiler toolchain (GCC, instead of LLVM for the Wasm binary), where the same nominal optimization level (O3) may have different effects. These results suggest that some work may still be required to completely fulfill Wasm’s promise of “near-native execution”. Nonetheless, the current performance is largely satisfactory for kana, and will likely improve over time as browser implementations evolve along with our understanding of the relevant optimizations.

**Table 2:** Analysis run times for several datasets with kana or a native executable. Values are reported in seconds with standard errors from 3 runs.

| Dataset | Number of cells | kana | Native |
| --- | --- | --- | --- |
| Zeisel | 3005 | $7.00 \pm 0.10$ | $5.60 \pm 0.05$ |
| Paul | 10368 | $17.59 \pm 0.20$ | $13.52 \pm 0.38$ |
| Bach | 25806 | $54.96 \pm 1.13$ | $43.33 \pm 0.39$ |
| Ernst | 68937 | $157.15 \pm 7.39$ | $114.67 \pm 1.86$ |
| Bacher | 104417 | $228.02 \pm 2.85$ | $170.32 \pm 1.34$ |
| Zilionis | 173954 | $272.265 \pm 4.22$ | $183.77 \pm 2.46$ |

#### 2.2.2 Parallelization with web workers

Web workers provide a simple mechanism for parallelization inside the browser. In kana, a web worker is used to run the compute-intensive analysis in its own thread so that the application’s main thread is free to respond to user interaction. We also use web workers to parallelize the operations within some analysis steps by compiling our C++ code with PThreads support; this instructs Emscripten to implement POSIX threads as web workers for transparent parallelization across genes or cells during calculation of QC metrics, nearest neighbor detection and marker scoring. Finally, we parallelize across steps by manually creating a separate web worker to execute the t-SNE, UMAP and clustering steps, which are independent of each other and can run concurrently.

A minor complication with PThreads is that we need to enable cross-origin isolation on the kana site. Briefly, this is a security measure that prevents inappropriate data access from malicious scripts in the same browsing context. Once cross-origin isolated, the site is permitted to use a SharedArrayBuffer for copy-free transfer of data between web workers. To achieve this, we need to serve the kana assets with the appropriate cross-origin headers – specifically, the embedder and opener policies. However, this may not always be possible, e.g., when hosting on institutional sites or public sites such as GitHub Pages. Instead, we use a service worker to cache and re-serve kana with the correct headers, ensuring that it can be easily deployed in a range of hosting environments.

As a matter of etiquette, we limit kana to two-thirds of the available threads at maximum utilization. The intention is to leave enough resources available for other activities to pass the time while waiting for the analysis to finish.

#### 2.2.3 Creating layered sparse matrices

To reduce memory usage for large single-cell count matrices, we use a “layered matrix” approach that splits the input matrix by row into 3 sparse submatrices. The first, second and third submatrices contain data for genes where all non-zero counts can fit into 8-bit, 16-bit and 32-bit unsigned integers, respectively. This improves memory efficiency for datasets with many cells but low sequencing coverage; most counts can be represented as 8-bit integers to save space, leaving a few high-abundance genes to be stored in the submatrices with larger integer types. A similar strategy is applied to row indices for non-zero elements in a sparse matrix, where most datasets have fewer than 60,000 genes and can be accommodated with 16-bit integers. This gives a theoretical usage of 3 bytes per non-zero element, e.g., a dataset with 30,000 genes and 100,000 cells at 5% density requires around 500 MB of memory in this data structure.

The layered matrix representation is implemented through the delayed binding mechanism in tatami. Specifically, we create the individual sparse submatrices and then create an abstract representation of the full matrix where the submatrices are combined by row. This preserves the memory-efficient representation while presenting an interface that mimics that of a single matrix. The layered representation can then be seamlessly used with all C++ code compatible with the tatami interface. (Note that the genes are permuted from their input order, which requires some extra attention in downstream analyses.)

#### 2.2.4 Examining memory usage

As an additional evaluation, we recorded the memory usage of the kana analyses (Table 3). For kana, we used the final size of the Wasm heap as a convenient proxy for the peak memory used by the application. This omits some memory usage on the JavaScript side but should capture the most memory-intensive data structures, particularly the count matrix. For comparison, we also recorded the maximum resident set size after each run of the native executable. In all cases, memory usage with Wasm was comparable to that of the native executable and maintained below 3 gigabytes, which is reasonable for a client device.

**Table 3:** Memory usage of kana or a native executable when analyzing datasets from Table 1. Values are reported in megabytes with standard errors from 3 runs. The number of non-zero elements for each dataset is also shown in millions.

| Dataset | Cells | Non-zeros | kana | Native |
| --- | --- | --- | --- | --- |
| Zeisel | 3005 | 11.3 | $148.31 \pm 0$ | $158.20 \pm 0.05$ |
| Paul | 10368 | 11.9 | $224.87 \pm 0$ | $306.26 \pm 0.06$ |
| Bach | 25806 | 61.2 | $761.12 \pm 0$ | $965.17 \pm 0.03$ |
| Ernst | 68937 | 163.1 | $1981.87 \pm 0$ | $2512.08 \pm 0.06$ |
| Bacher | 104417 | 143.0 | $2165.94 \pm 0$ | $2437.75 \pm 1.34$ |
| Zilionis | 173954 | 53.7 | $2480.81 \pm 0$ | $2600.66 \pm 0.04$ |

Importantly, kana’s memory usage lies below Wasm’s hard limit of 4 gigabytes of addressable memory. This limit is a consequence of the use of 32-bit pointers in the current Wasm specification, and exceeding this limit will cause allocation errors and application failure. Future Wasm releases may support 64-bit pointers [61] or multiple memories [57]; if these proposals are accepted, kana will be able to process datasets beyond the 4 GB limit.

### 2.3 User interface concepts

#### 2.3.1 Interlinked graphics

Different panels of the kana application (Figure 2) can share information with each other to facilitate interactive exploration of the dataset and analysis results. For example, we can color the embeddings according to the expression of a gene selected from the marker table. Upon selection, kana retrieves the log-normalized expression values for that gene from the sparse matrix and passes this data to the scatter plot for coloring. Users can also adjust the color gradient to improve contrast and highlight differences at particular ranges of expression.

A more complex example involves the detection of new marker genes for a custom selection of cells. Users can create a custom selection by brushing on regions of interest in the embedding panel. If this selection is saved, kana will perform a differential expression analysis to detect upregulation inside the selection compared to all other cells. Statistics for each gene are then shown in the marker table for examination, with an additional option to perform gene set enrichment analyses on the top markers. Each custom selection and its statistics are treated as part of the analysis state and are saved during export.

Inspired by the gallery in Cirrocumulus [20], users can save specific views of the embeddings for later perusal. This is useful for simultaneously viewing multiple plots colored by different marker genes to identify cell populations.

#### 2.3.2 Progressive rendering

Each result is immediately rendered on the interface once the corresponding analysis step is complete. For example, the distribution of QC metrics appear once the QC step is finished, followed by the plot of the percentage of variance explained once the PCA is done, and so on. This improves application performance by avoiding the rendering bottleneck that would otherwise occur if all plots were drawn at once. It also serves as a visual progress indicator and improves the user experience by showing meaningful content as soon as possible. The marker table is a more subtle example of progressive rendering. Only the genes in the current view of the table are rendered, with the remaining visual elements being dynamically created as the user scrolls up or down to see other genes in the ranking. This ensures that we do not waste time rendering tens of thousands of rows when only the top few are likely to be viewed.

Given that kana is computing the embeddings in the background, we can actively monitor the change in the coordinates for each cell across t-SNE iterations or UMAP epochs. We do so by extracting the coordinates at regular intervals and rendering them on the embedding panel with WebGL, effectively creating an animation of the embedding as it is refined over time. This is mostly present for entertainment value but may provide some educational insights into the process of creating the embeddings (e.g., early exaggeration for t-SNE).

#### 2.3.3 Exploration mode

kana provides an “exploration mode” for datasets that contain pre-computed results. In this mode, many of the analysis steps are skipped as their results are already available inside the dataset file. For example, we can extract the reduced dimensions from the “reducedDims” slot of a SingleCellExperiment contained inside an RDS file, or from the “obsm” group of a H5AD file. Similarly, we can extract clustering results from the column annotations of these representations. This functionality allows users to explore analysis results using the kana’s interface without waiting for an unnecessary re-analysis. The exceptions are that of marker detection and gene set enrichment, which are computed on demand based on user interaction; this is acceptable as these steps are relatively cheap and their results are often absent from many representations.

#### 2.3.4 Exporting and reloading

Users save their analysis configuration to the browser’s cache via the IndexedDB system. This dumps the analysis parameters into a JSON file inside the cache, along with the input data files. kana will attempt to deduplicate input files in the cache based on the file name, size and MD5 checksum. This reduces disk usage when storing multiple analysis states for the same dataset. Users can then reload their saved configurations in a new browser session with a single click.

Alternatively, users may export the analysis parameters and input files into a Zip archive that is downloaded to their machine. Supplying this file to kana will then restore the previous analysis in the same manner as using the browser cache. The benefit of this approach is that files exported by one user can be reloaded by other users on different machines, allowing users to easily share their configurations with each other by simply transferring the Zip file.

Finally, users can export the analysis results into the ArtifactDB format (https://github.com/ArtifactDB). Briefly, this creates a language-agnostic representation of Bioconductor’s SingleCellExperiment object using a combination of HDF5 and CSV files. This representation can be loaded into the user’s programming framework of choice for further analyses with other tools. For example, the alabaster Bioconductor package [49] will reconstitute a SingleCellExperiment from the kana-exported files. The same files can also be imported into kana’s exploration mode for continued inspection in the browser.

## 3 Discussion

Single-cell data analysis on the client is not a particularly novel concept. In many cases, the dataset already lives on the user’s machine, so this is a natural place to perform the analysis with the tool of choice (typically R or Python). kana’s key innovation lies in its use of modern web technologies to perform the analysis directly in the browser. This eliminates the difficulties of software installation and makes the analysis accessible to a non-programming audience. At the same time, we retain all the benefits of client-side operations. Specifically, our availability only depends on the client device, not on a backend server; there is no latency from transferring data or results across a network; ownership of the dataset clearly remains with the user, avoiding issues with data privacy and associated regulations; and compute time is cheap, if not effectively free.

Client-side compute has interesting scalability characteristics compared to a traditional backend approach. Most obviously, we are constrained by the computational resources available on the client machine, which can vary from modest (e.g., most laptops) to extremely basic (e.g., mobile devices). This limits the size of any single dataset that can be analyzed by a particular client. However, in other respects, client-side compute is more scalable than backend compute; the former automatically distributes analyses of many datasets across any number of machines at no cost and with no configuration. This is especially relevant for web applications like kana where the maintainers would otherwise be responsible for provisioning backend computing resources. The cloud might be infinitely scalable but - unfortunately - our bank accounts are not.

That said, how do we deal with large datasets? This is not straightforward with the web technologies currently used in kana - our available memory is limited by the Wasm specification and our CPU utilization is (somewhat artificially) capped, so even if a powerful client device was available, kana would not be able to fully exploit its capabilities. Fortunately, our use of C++ means that we are not limited to computation in the browser. We can easily provide wrappers to the same underlying libraries in any client-side framework, e.g., as a command-line tool or as an extension to existing data science ecosystems. For example, the scran.chan package [48] allows users to repeat the kana calculations in an R session, where there are no restrictions on memory usage or thread utilization. The same approach can be used for other languages that support a foreign function interface like Python or Julia. Indeed, one could use the wrapped C++ libraries to run large analyses on a sufficiently provisioned backend, export the results in a kana-compatible file format, and then serve the exported results to client machines for exploration in the browser.

Future work will involve adapting kana to take advantage of the continued evolution of web technologies. This includes further optimizations to the Wasm specification and its implementations, as well as upcoming standards like WebGPU to leverage graphics hardware for general computation. By incorporating these advances, applications like kana can extend the capabilities of the browser for compute-intensive bioinformatics in the client.

## 4 Conclusions

kana leverages WebAssembly and other web technologies to efficiently analyze single-cell ‘omics data in the browser. This empowers all scientists to perform single-cell analyses without the need to install any additional software or learn any programming language. By distributing the analyses to the client devices, kana avoids the provisioning and data transfer problems associated with a back-end server. This paradigm of client-side compute provides certain cost and scalability benefits that may be generally beneficial to other web applications that are commonly used by the bioinformatics community.

## 5 Methods

Here we summarize the key software repositories used in this paper. For brevity, we will omit the various C++ libraries that have been listed previously.

- The kana application is deployed at https://www.kanaverse.org/kana with source code at https://github.com/kanaverse/kana.
- The bakana library is available at https://npmjs.com/package/bakana with source code at https://github.com/kanaverse/bakana.
- The scran.js library is available at https://npmjs.com/package/scran.js with source code at https://github.com/kanaverse/scran.js.
- The scran.chan R package based on the same C++ libraries is available at https://github.com/LTLA/scran.chan.
- The command-line interface based on the same C++ libraries is available at https://github.com/LTLA/scran-cli.
- All datasets used here can be downloaded from https://github.com/kanaverse/random-test-files.
- Benchmarking code and results are available at https://github.com/kanaverse/kana.perf.

## 6 Declarations

### 6.1 Ethics approval

Not applicable.

### 6.2 Consent for publication

All authors consent to the publication of this manuscript.

### 6.3 Availability of data and materials

All datasets are publicly available as described in Table 1. All software is available at the URLs listed in the Methods section.

### 6.4 Competing interesets

The authors declare that they have no competing interests.

### 6.6 Author contributions

Both authors were involved in the conception and design of this application. AL developed the C++ libraries and most of the Javascript bindings. JK wrote the kana application itself, including the user interface and the various visual components. Both authors were involved in the writing of this manuscript.

## 6.7 Acknowledgements

We thank Michael Lawrence, Hector Corrada Bravo and Adrian Waddell for their suggestions to improve this manuscript.

